# Stable neutralizing antibody levels six months after mild and severe COVID-19 episode

**DOI:** 10.1101/2020.11.22.389056

**Authors:** Edwards Pradenas, Benjamin Trinité, Víctor Urrea, Silvia Marfil, Carlos Ávila-Nieto, María Luisa Rodríguez de la Concepción, Ferran Tarrés-Freixas, Silvia Pérez-Yanes, Carla Rovirosa, Erola Ainsua-Enrich, Jordi Rodon, Júlia Vergara-Alert, Joaquim Segalés, Victor Guallar, Alfonso Valencia, Nuria Izquierdo-Useros, Roger Paredes, Lourdes Mateu, Anna Chamorro, Marta Massanella, Jorge Carrillo, Bonaventura Clotet, Julià Blanco

**Author notes:** Corresponding author: Julià Blanco, PhD Senior Researcher, Institut de Recerca de la Sida. IrsiCaixa, IGTP, Hospital Germans Trias i Pujol Ctra. de Canyet s/n. 2a Planta Maternal. 08916 Badalona. Barcelona. Equal contribution (Edwards Pradenas, Benjamin Trinité).

## Abstract

Understanding mid-term kinetics of immunity to SARS-CoV-2 is the cornerstone for public health control of the pandemic and vaccine development. However, current evidence is rather based on limited measurements, thus losing sight of the temporal pattern of these changes^1–6^. In this longitudinal analysis, conducted on a prospective cohort of COVID-19 patients followed up to 242 days, we found that individuals with mild or asymptomatic infection experienced an insignificant decay in neutralizing activity that persisted six months after symptom onset or diagnosis. Hospitalized individuals showed higher neutralizing titers, which decreased following a two-phase pattern, with an initial rapid decline that significantly slowed after day 80. Despite this initial decay, neutralizing activity at six months remained higher among hospitalized individuals. The slow decline in neutralizing activity at mid-term contrasted with the steep slope of antibody titers change, reinforcing the hypothesis that the quality of immune response evolves over the post-convalescent stage^4,5^.

---

Our analysis included 210 patients with RT-PCR-confirmed SARS-CoV-2 infection, recruited during the first and second waves of the COVID-19 epidemic in Catalonia (North-East Spain). Of them, 106 (50.5%) had a mild or asymptomatic infection, and 104 (49.5%) required hospitalization because of respiratory compromise (Table 1). We collected samples periodically throughout a maximum follow-up period of 242 days (Figure S1, Supplementary Information). Most study participants developed a neutralizing humoral response against SARS-CoV-2 HIV-based pseudoviruses, that was confirmed using infectious viruses. However, in line with trends reported elsewhere^2,3^, mildly affected or asymptomatic individuals developed a 10-fold lower maximal neutralization titer than those who required hospitalization when the full dataset was analyzed (*p*<0.0001, Mann-Whitney test; Fig. 1a). The higher number of determinations obtained from hospitalized individuals during the acute phase permitted the clear observation of a sharp initial response (Fig. 1b-c), also reported in previous analyses of the early response^7–11^. This was visible for individuals recruited during both the first (March-June 2020) and the second (July-October 2020) waves of COVID-19 pandemic in Catalonia. A longitudinal analysis fitted to a four-parameter logistic model of increase defined a 30-day sharpening phase after symptom onset, irrespective of the wave in which hospital admission occurred. Half maximal neutralization activity was achieved on day 10 (95% confidence interval, CI 8-11); 80% maximal response, which corresponded to 3.97 logs (i.e., 9,333 reciprocal dilution), was achieved on day 14 (Fig. 1d). Based on this finding, we assumed no significant differences between the two waves regarding early neutralizing response and we decided to set day 30 after symptom onset as a starting point for the longitudinal analysis of immune response at the mid-term.

**Fig 1.**
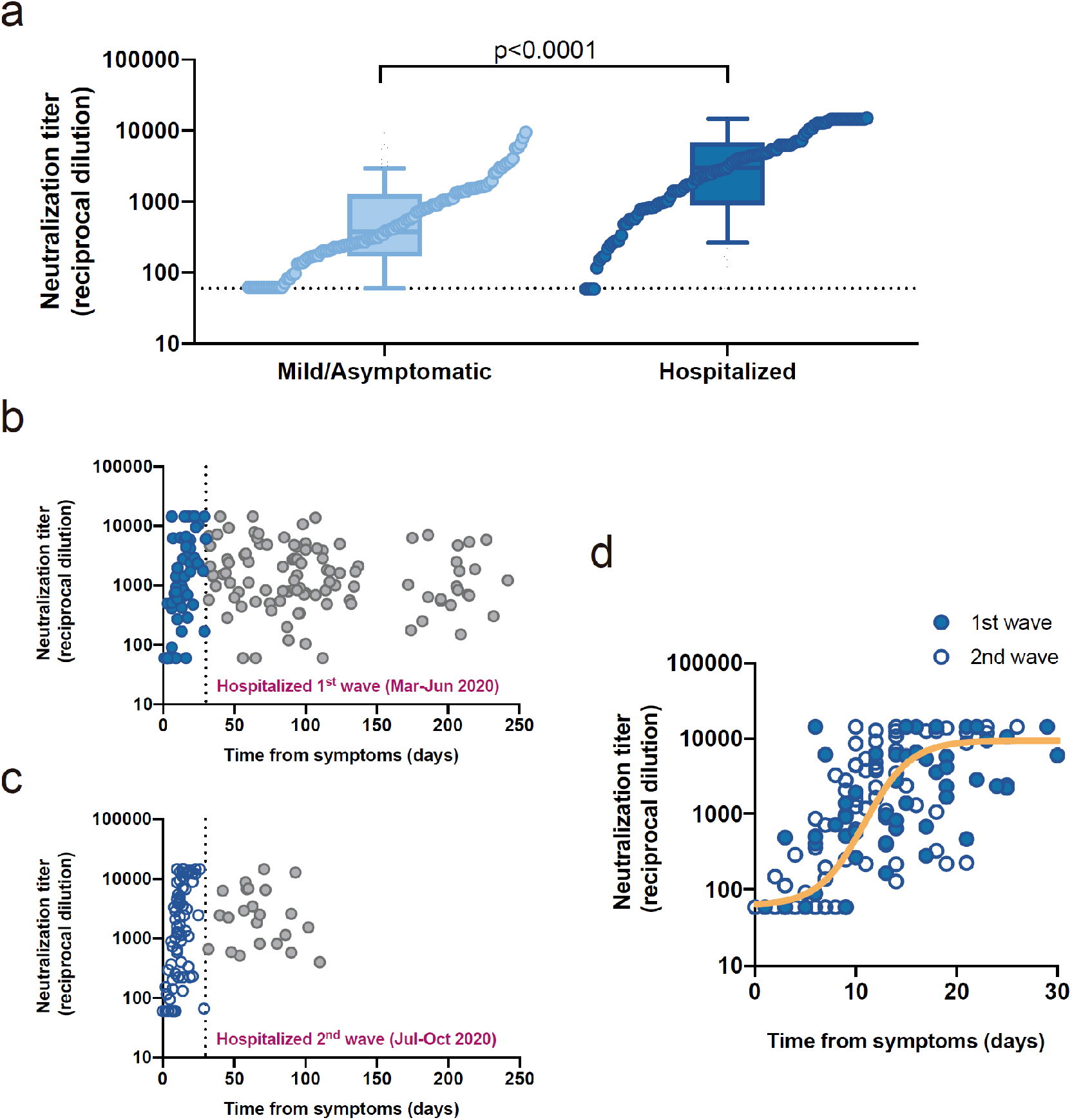
Neutralizing activity among study participants. **a**, Maximal neutralization titer of 210 individuals recruited, according to disease severity (light and dark blue for mild/asymptomatic and hospitalized individuals, respectively). Boxes show the median and the interquartile range, and bars the 10^th^ and 90^th^ percentiles. Distributions were compared using the Mann-Whitney test. Individual values are ranked for comparative purposes. **b** and **c**, Longitudinal dot plot of neutralizing activity among hospitalized individuals admitted during the first (**b**) and second (**c**) waves of the COVID-19 epidemic in our area; filled (**b**) and empty (**c**) blue dots show the early (i.e., 30 days after diagnosis) increasing phase. **d**, Magnification of the early phase for individuals admitted during the first (filled symbols) and second (empty symbols) waves. No differences between waves were observed. The solid line shows the non-linear fit (mixed-model estimate) for the whole dataset (125 samples, 55 individuals analyzed). Two samples from late seroconverters (one from each wave, grey dots) were excluded from the analysis.

**Table 1.** Characteristics of individuals included in analysis

|  | Mild/asymptomatic<br>(n=106) | Hospitalized<br>(n=104) | *p value* |
| --- | --- | --- | --- |
| Gender (female), *n (%)* | 72 (68) | 46 (44) | 0.0006<sup>a</sup> |
| Age (years), *median [IQR]* | 46.5 [38 - 54] | 57.5 [46 - 66] | <0.0001<sup>b</sup> |
| Individuals with 2 or more samples, *n (%)* | 52 (49) | 59 (57) | 0.278<sup>a</sup> |
| Wave of COVID-19 outbreak (first), *n (%)* | 96 (91) | 73 (70) |  |
| Severity, *n (%)* |  |  |  |
| Asymptomatic | 8 (8) | --- |  |
| Mild | 98 (92) | --- |  |
| Hospitalized non-severe | --- | 59 (56.7) |  |
| Hospitalized severe | --- | 37 (35.6) |  |
| Hospitalized (intensive care unit) | --- | 8 (7.7) |  |

IQR: interquartile range (25th and 75th percentiles), <sup>a</sup> Chi square test, <sup>b</sup> Mann-Whiney test

While the early humoral response after SARS-CoV-2 infection has been thoroughly described^7–11^, current data on the decay of antibody levels beyond the convalescent stage depict a heterogeneous scenario with limited information on the neutralizing activity throughout the follow-up period^1–3^. Various authors have recently suggested more complex kinetics of neutralizing activity decay, with clonotype-, epitope-, or subject-specific patterns that evolve in terms of potency and resistance to epitope mutations^4,5,12^. The longitudinal modeling of the neutralizing activity at mid-term in our cohort revealed a nearly flat slope (i.e., not significantly different from 0, with half-life 2134 days) in individuals with asymptomatic infection or mild disease (Fig. 2a). Conversely, the decrease of neutralizing activity in hospitalized individuals showed a two-phase pattern, with a rapid decay (half-life 31 days) until day 80 that slowed down to a flat slope (half-life 753 days) from that time point on (Fig. 2b).

**Fig 2.**
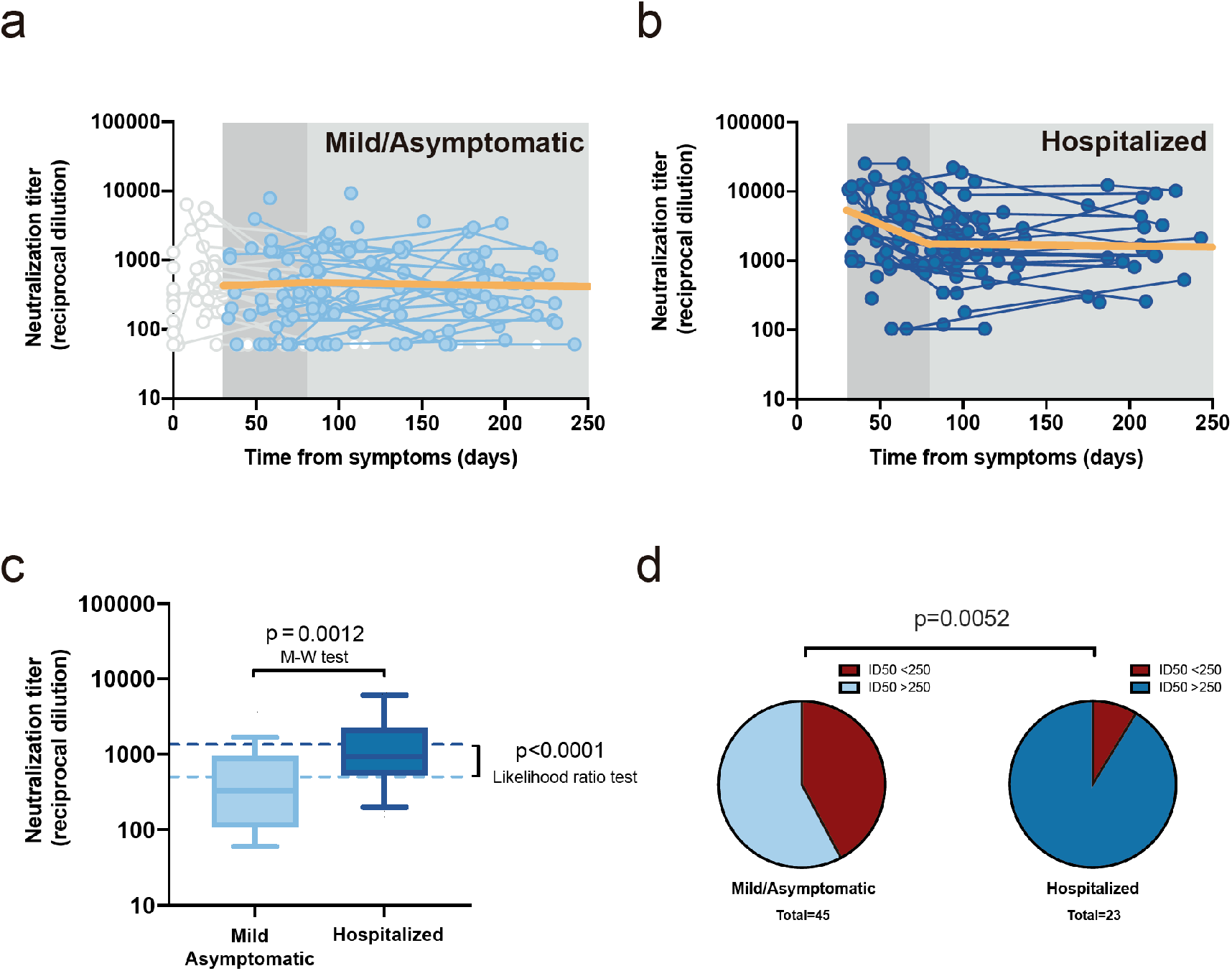
Longitudinal analysis of neutralizing activity. **a**, Individual measurements (dots) and linear mixed model (solid orange line) of the longitudinal analysis for mild or asymptomatic individuals beyond day 30 (single-phase slope −0.00014; *p*=0.75, likelihood ratio test; estimated half-life 2,134 days). Time points preceding day 30 as well as participants only showing undetectable titers were excluded from the analysis, values are shown but grayed out. **b**, the corresponding analysis for hospitalized individuals (the slopes of the linear fit for the first and second phase were -0.0096 [*p*=0.0002] [half-life 31 days] and −00004 [half-life 753 days] [*p*=0.78], respectively). **c**, Distribution of neutralizing activity six months after infection in both disease severity groups. Experimental values of mean neutralizing activities in the period 135-to-242 days as summarized in box-plots (as in Figure 1a; Mann-Whitney test for comparative analysis) and modeled data as dotted lines (likelihood ratio test for comparative analysis). **d**, Frequency of long-term neutralizers (i.e., individuals with mean neutralizing activity >250 in the 135-242 days period) in each severity subgroup (Chi square test p value is shown).

The characterization of the neutralizing activity behavior at mid-term should ultimately project the proportion of post-convalescent individuals protected against new infections in the mid- and long-term. The limited number of measures and lack of a clear threshold of neutralizing activity for preventing SARS-CoV-2 infection precluded assessing this outcome using survival analysis. Alternatively, we explored the neutralizing activity at the end of our 6-month follow-up period. Based on the mixed-effects model obtained from the longitudinal analysis, we estimated a stable mid-term neutralizing activity of 2.72 and 3.16 log for the mild/asymptomatic and hospitalized subgroups, respectively (*p*<0.0001; likelihood ratio test, Fig. 2c, dotted lines). This estimate was consistent with the observed values for the last measurement taken between days 135 and 242, a time frame centered on day 180 (Fig. 2c, box plots). Likewise, the value distribution at this time frame showed significant differences between mild/asymptomatic (median 2.5; IQR 2.0–3.0) and hospitalized (3.0; 2.7–3.3) individuals (*p*=0.0012, Mann-Whitney test). To date, no clear cut-off for a neutralizing activity that protects against new reinfection has been established. Nevertheless, data gathered from high attack rate events suggest that neutralizing activities between 1:161 and 1:3,082 are strong enough to prevent infection^13^. Hence, we assumed that reinfections would be unlikely among individuals above the 1:250 cut-off. Of the 23 hospitalized individuals with measurement beyond day 135, 21 (91%) had a mean neutralizing activity value above 1:250 in the 135-242 days period and were thus considered long-term neutralizers. The corresponding proportion in the mild/asymptomatic group (42%; 19/45) was significantly lower (*p*=0.0052, Chi exact test, Fig. 2d). Although this number must be taken cautiously due to the cut-off assumption, our finding suggests that hospitalized patients have a higher capacity for long-term neutralization, despite the faster initial decay in neutralization activity.

It has recently been proposed that the kinetics of neutralizing activity may not mirror those of antibody titers^4^. Hence, we investigated the change in serum titers of IgG in a subset of 28 individuals (14 in each severity group) with the most extended follow-up period. The analysis included antibodies against the receptor-binding domain (RBD) and S2 subunit of the S protein, both associated with potential neutralizing activity; and the nucleoprotein (NP), which are very abundant, albeit unable to neutralize the SARS-CoV-2 directly^14^. The longitudinal analysis revealed a one-phase significant (*p*<0.0001) steady decay pattern of the three tested antibodies, which was notably faster in IgG anti-NP (Fig. 3a-c). The half-life of anti-RBD, anti-S2, and anti-NP antibodies for the period beyond day 30 were 86, 108, and 59 days, respectively. These values were consistent with those reported by Wheatley et al., estimated on a 160-day time frame^4^. Although the limited sample size of this sub-analysis precluded independent modeling of the decay in mild/asymptomatic and hospitalized patients, the latter showed significantly higher titers of anti-S2 at the end of the follow-up period (Figure S2), whereas no significant differences were found in other antibodies regarding disease status. Interestingly, in this subset of individuals, the decay in antibody titers contrasted with the behavior of neutralizing activity, which fitted to a two-phase model―as in the whole dataset―with a rapid decay until day 80 (slope 0.014, half-life 22 days) and a flat slope (i.e., not significantly different from 0) afterward (Fig. 3d).

**Fig 3.**
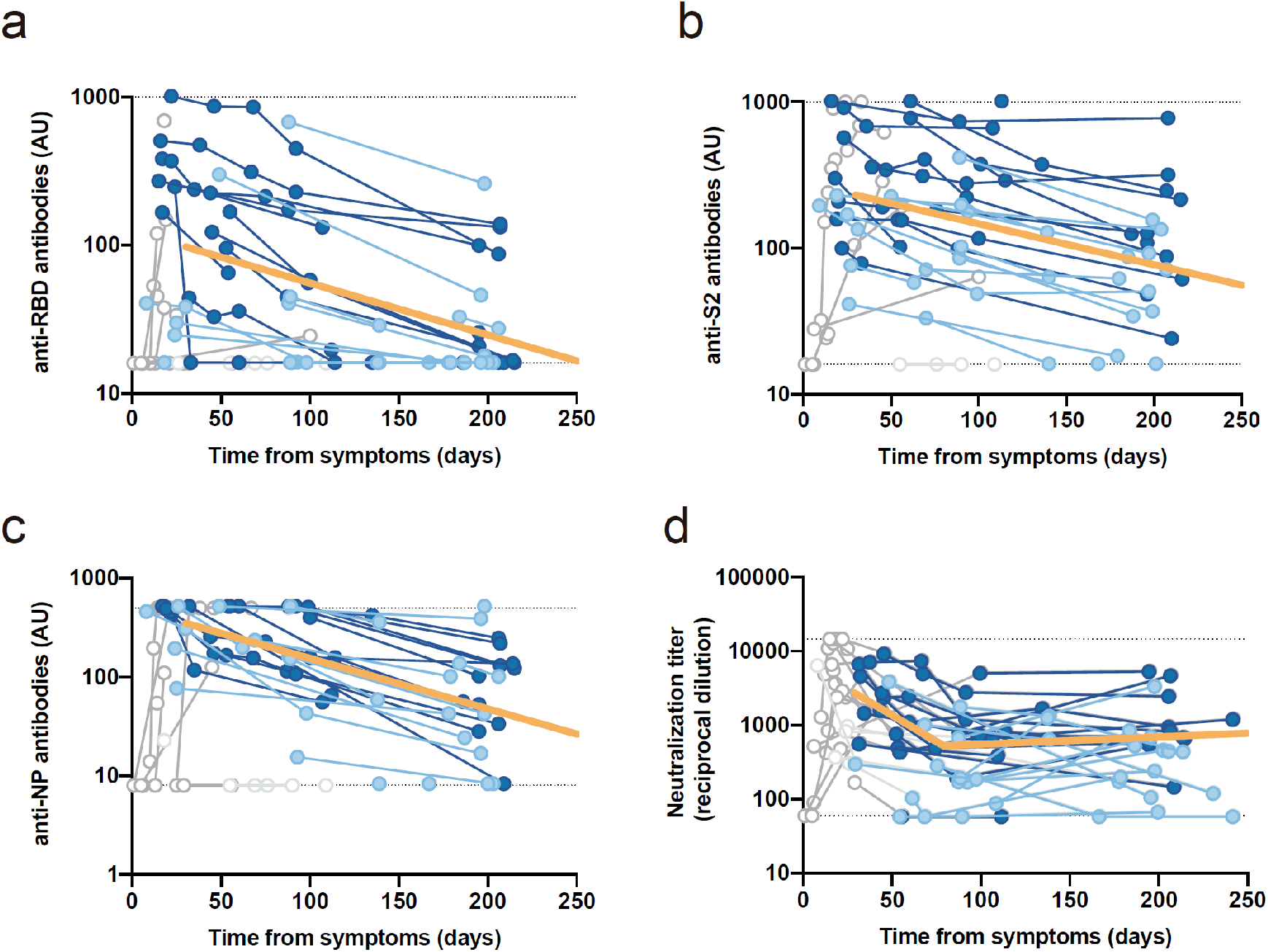
Longitudinal analysis of IgG titers. **a**, anti-receptor binding domain (RBD). **b,** anti-S2. **c,** anti-nucleoprotein. **d,** overall neutralizing activity in the same set of samples. All analyses were performed on a subset of individuals with largest follow-up (n=14 for mild/asymptomatic in light blue and n=14 for hospitalized in dark blue; total no. of samples 94). Solid orange lines show the linear mixed model estimate for the period beyond day 30. Kinetics of antibody decay (panels a-c) were calculated excluding timepoints preceding the maximal values for each patient. Kinetics of neutralizing antibodies excluded samples preceding day 30 (as in Fig 2a/b). All excluded values are shown but grayed out.

Complementary data on the binding affinity and B-cell clone abundance at the same time points would provide a more comprehensive picture to explain this divergent trend. However, our findings support the hypothesis of Gaebler et al., who suggested that the accumulation of IgG somatic mutations―and subsequent production of antibodies with increased neutralizing potency―allow the maintenance of neutralizing activity levels, despite the decline in specific antibody titers^5^. Of note, our follow-up period encompassed two waves of the COVID-19 outbreak in our country. Individuals infected during the first wave were likely to be exposed to high viral pressure in their environment, potentially favoring further virus exposure that may also contribute to boosting humoral responses, adding to the mechanism proposed by Gaebler et al^5^.

Our analysis is limited by the reduced sample size, particularly in the mild/asymptomatic subgroup, for which we failed to identify a two-phase pattern decay of neutralizing activity. Despite the limited sample size, the availability of multiple measures along the follow-up period allowed us to provide a longitudinal perspective on neutralizing activity, and antibody titer behavior. This analysis supplements current evidence regarding mid-term immunity against SARS-CoV-2, based on two- or three-point measurements^3–5^. Our longitudinal analysis confirmed the slow decay and mid-term maintenance of neutralizing activity observed in other cohorts with a 5-to-11% prevalence of hospitalized patients^3,5^. In this regard, the two-phase behavioral pattern of neutralizing activity observed in hospitalized individuals suggests that the rapid decay reported in previous characterizations^1^ might be due to the abundance of individuals in this early phase. Furthermore, apparent inconsistencies found between the declines of neutralizing activity and IgG titers reinforce the idea proposed by other authors that the behavior of antibody titers may not mirror the neutralizing activity^4^. Taken together, current evidence on immunity to SARS-CoV-2 infection suggests stability of neutralizing activity, pointing towards an optimistic scenario for the establishment of infection- or vaccine-mediated herd immunity. Still, long-term data available on other human coronaviruses show waning of antibodies 1-to-2 years after infection^15,16^, with uncertainty regarding the immune response behavior in the context of vaccine-mediated immunity^17^. The continuity of our prospective cohort of individuals recovered from SARS-CoV-2 infection will provide novel insights into the long-term kinetics of the immune response.

## Methods

### Study overview and subjects

The study was approved by the Hospital Ethics Committee Board from Hospital Universitari Germans Trias i Pujol, HUGTiP (reference PI-20-122 and PI-20-217) and all participants provided written informed consent before inclusion.

Plasma samples were obtained from individuals of the prospective KING cohort of the HUGTiP (Badalona, Spain), designed to characterize virological and immunological features of SARS-CoV-2 infection. The recruitment period lasted from March to October 2020, thus covering the first and second wave of COVID-19 outbreak in Catalonia (dadescovid.cat). The KING cohort included more than 200 individuals with a documented positive RT-qPCR result from nasopharyngeal swab and/or a positive serological ELISA test performed in our hospital, irrespective of age and disease severity―including asymptomatic status. Individuals were recruited in various settings, including primary care, hospital, and epidemiological surveillance based on contact tracing. We collected plasma samples at the time of COVID-19 diagnosis and at 3 and 6 months. Additionally, hospitalized individuals were sampled twice a week during the acute phase. For the longitudinal analysis of immune response kinetics, we selected only samples from patients with at least two determinations during the follow-up period (30-242 days).

### Humoral response determination by Enzyme-Linked Immunosorbent Assay (ELISA)

The humoral response against SARS-CoV-2 was evaluated using an in-house sandwich-ELISA. The following SARS-CoV-2 antigens were purchased from Sino Biological (Eschborn, Germany): S2 (Ser686-Pro1213, produced using insect cells), RBD (Arg319-Phe541, produced in HEK293 cells), and nucleocapsid protein (NP) (whole molecule, produced in insect cells). Briefly, Nunc MaxiSorp ELISA plates were coated with 50 μL of capture antibody (anti-6x-His antibody clone HIS.H8; 2 μg/mL; Thermo Fisher Scientific, Waltham, MA, USA) in phosphate-buffered saline (PBS) overnight at 4°C. After washing, plates were blocked with 1% BSA in PBS solution (Miltenyi Biotec, Bergisch Gladbach, Germany) for two hours at room temperature. Plates were rewashed, and the antigens were added at 1 μg/mL concentration (50 μL/well) and incubated overnight at 4°C. Plasma samples were heat-inactivated before use (56°C for 30 minutes) and analyzed by duplicate with each SARS-CoV-2 antigen in the same plate, along with antigen-free wells to assess background. Serial dilution of a positive plasma sample was used as standard. A pool of ten plasmas collected from healthy controls before August 2019 was used as a negative control. Standards, negative control, and plasma samples were diluted in blocking buffer. After blocking, the plates were washed and standards, negative control, and diluted plasma samples were added to the plate (50 μL/well) and incubated for one hour at room temperature. The HRP conjugated (Fab)2 goat anti-human IgG (Fc specific) (1/20,000) (Jackson ImmunoResearch, Ely, UK) was used as a secondary antibody and incubated for 30 minutes at room temperature. Plates were revealed with o-Phenylenediamine dihydrochloride (OPD) (Sigma-Aldrich, St. Louis, MO, USA) and the reaction was stopped using 4N of H_2_SO_4_ (Sigma-Aldrich). The signal was analyzed as the optical density (OD) at 492 nm with noise correction at 620 nm. The specific signal for each antigen was calculated after subtracting the background signal obtained for each sample in antigen-free wells. Results were calculated using a 5 Parameter Logistic (5PL) curve and expressed as arbitrary units (AU) according to the standard.

### Pseudovirus generation and neutralization assay

HIV reporter pseudoviruses expressing SARS-CoV-2 S protein and Luciferase were generated. pNL4-3.Luc.R-.E- was obtained from the NIH AIDS Reagent Program^18^. SARS-CoV-2.SctΔ19 was generated (GeneArt) from the full protein sequence of SARS-CoV-2 spike with a deletion of the last 19 amino acids in C-terminal^19^, human-codon optimized and inserted into pcDNA3.4-TOPO. Expi293F cells were transfected using ExpiFectamine 293 Reagent (Thermo Fisher Scientific) with pNL4-3.Luc.R-.E- and SARS-CoV-2.SctΔ19 at a 24:1 ratio, respectively. Control pseudoviruses were obtained by replacing the S protein expression plasmid with a VSV-G protein expression plasmid as reported previously^20^. Supernatants were harvested 48 hours after transfection, filtered at 0.45 μm, frozen, and titrated on HEK293T cells overexpressing WT human ACE-2 (Integral Molecular, Philadelphia, PA, USA).

Neutralization assays were performed in duplicate. Briefly, in Nunc 96-well cell culture plates (Thermo Fisher Scientific), 200 TCID_50_ of pseudovirus was preincubated with three-fold serial dilutions (1/60 – 1/14,580) of the heat-inactivated plasma samples (see above) for 1 hour at 37°C. Then, 2×10^4^ HEK293T/hACE2 cells (Integral Molecular) treated with DEAE-Dextran (Sigma-Aldrich) were added. Results were read after 48 hours using the EnSight Multimode Plate Reader and BriteLite Plus Luciferase reagent (PerkinElmer, Waltham, MA, USA). The values were normalized, and the ID_50_ was calculated by plotting the log of plasma dilution vs. response―Variable slope (four parameters) in Prism 8.4.3 (GraphPad Software, San Diego, CA, USA). ID_50_ was defined as the reciprocal of the dilution that inhibited 50% of the infection.

### Statistical analysis

Continuous variables were described using medians and the interquartile range (IQR, defined by the 25^th^ and 75^th^ percentiles), whereas categorical factors were reported as percentages over available data. Quantitative variables between severity groups were compared using the Mann-Whitney test, and percentages using the chi-squared test. Kinetics of neutralizing activity and antibody titers were estimated from symptom onset―or serological diagnosis in asymptomatic individuals―and modeled using mixed-effects models and in two steps. First, a 4-parameter logistic function was adjusted for the first 30 days after diagnosis using non-linear mixed models. Mid-term decay was analyzed using a piecewise regression with two decline slopes for data beyond 30 days, with a breakpoint at 80 days. For the latter analysis, linear mixed-effect models with random intercepts and slopes were used, and different breakpoints were tested; the one with the best adjustment was chosen. For the longitudinal analysis of neutralizing activity, patients were grouped into two severity groups according to the WHO progression scale as proposed elsewhere^21^: asymptomatic or mild (levels 1-3), and hospitalized (levels 4-10). Differences between the two severity groups were assessed using the likelihood ratio test. The longitudinal analysis of antibody titers was performed on a subset of 28 individuals (14 in each severity group) with the highest number of measures during the follow-up; owing to the limited sample size, all 28 individuals were analyzed as a single group. All analyses were performed with Prism 8.4.3 (GraphPad Software) and R version 4.0 (R Foundation for Statistical Computing). Mixed-effects models was fitted using “nlme” R package.

## Acknowledgements

This work was partially funded by Grifols, the *Departament de Salut* of the *Generalitat de Catalunya* (grant DSL0016 to JB and Grant DSL015 to JC), the Spanish Health Institute *Carlos III* (Grant PI17/01518 and PI18/01332 to JC) and the crowdfunding initiatives #joemcorono, BonPreu/Esclat and Correos. The funders had no role in study design, data collection and analysis, the decision to publish, or the preparation of the manuscript. EP was supported by a doctoral grant from National Agency for Research and Development of Chile (ANID): Grant 72180406. CA-N was supported by a doctoral grant from Secretaria d’Universitats i Recerca de la Generalitat de Catalunya i del Fons Social Europeu (FI). SP-Y is supported by Fundación Canaria Doctor Manuel Morales and by a doctoral grant from Universidad de La Laguna.

We are deeply grateful to all participants and the technical staff of IrsiCaixa for sample processing. Gerard Carot-Sans provided medical writing support during the preparation of the manuscript.

## Author contributions

JB and BC designed and coordinated the study. EP, BT, SM, CA-N, MLR, FT-F, SP-Y, CR, EA-E, JR, JV-A, JS and NI-U performed and analyzed pseudovirus production, neutralization and ELISA assays. VU performed statistical analysis. RP, LM, AC, MM, VG, AV and JC selected patients and coordinated data. JB and Gerard Carot-Sans have drafted the first version of the manuscript and all authors have made substantial contributions to the revision of the subsequent versions. All authors have approved the submitted version of the manuscript and have agreed both to be personally accountable for the author’s own contributions and to ensure that questions related to the accuracy or integrity of any part of the work.

