## Supplementary Information for "Stable neutralizing antibody levels six months after mild and severe COVID-19 episode"

[1]IrsiCaixa AIDS Research Institute, Germans Trias i Pujol Research Institute (IGTP), Can Ruti Campus, UAB, 08916, Badalona, Catalonia, Spain
[2]Laboratorio de Inmunología Celular y Viral, Unidad de Farmacología, Sección de Medicina, Facultad de Ciencias de la Salud, Universidad de La Laguna (ULL), La Laguna, 38071, Tenerife, Spain.
[3]IRTA Centre de Recerca en Sanitat Animal (CReSA, IRTA-UAB), Campus de la UAB, 08193, Bellaterra, Catalonia, Spain
[4]UAB, Centre de Recerca en Sanitat Animal (IRTA-UAB), Campus de la UAB, 08193, Bellaterra, Catalonia, Spain
[5]Departament de Sanitat i Anatomia Animals, Facultat de Veterinària, UAB, 08193, Bellaterra, Catalonia, Spain
[6]Barcelona Supercomputing Center, 08034, Barcelona, Catalonia, Spain
[7]Catalan Institution for Research and Advanced Studies (ICREA), 08010, Barcelona, Catalonia, Spain
[8]Infectious Diseases Department, Fight against AIDS Foundation (FLS), Germans Trias i Pujol Hospital, 08916, Badalona, Catalonia, Spain
[9]University of Vic–Central University of Catalonia (UVic-UCC), 08500, Vic, Catalonia, Spain
[¶]Equal contribution

***Corresponding author:**

Julià Blanco, PhD
Senior Researcher
Institut de Recerca de la Sida. IrsiCaixa, IGTP, Hospital Germans Trias i Pujol
Ctra. de Canyet s/n. 2a Planta Maternal. 08916 Badalona. Barcelona


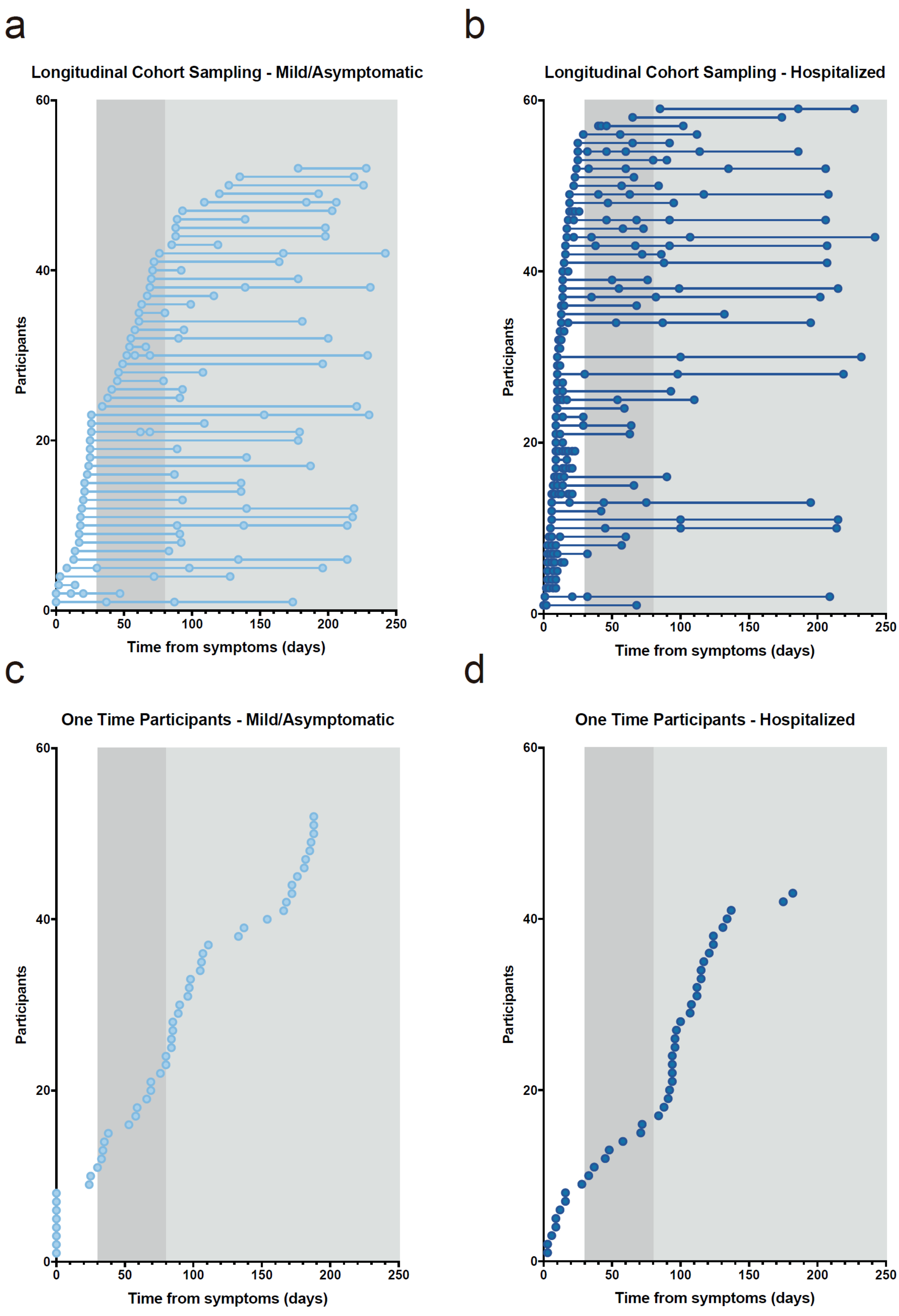


**Figure S1. Patient and sampling distribution across the follow-up period.** Top panels show the time points for sample collection among mild/asymptomatic (**a**) and hospitalized (**b**) individuals. Bottom panels show the time points for samples of individuals with a single measurement: **c**, mild/asymptomatic; **d**, hospitalized. Time count starts on the day of symptom onset, except for asymptomatic individuals, for whom the serological diagnosis was considered. The areas define the periods considered for the longitudinal analysis: days 0-30 (white), 30-80 (dark grey) and after 80 days (light grey).

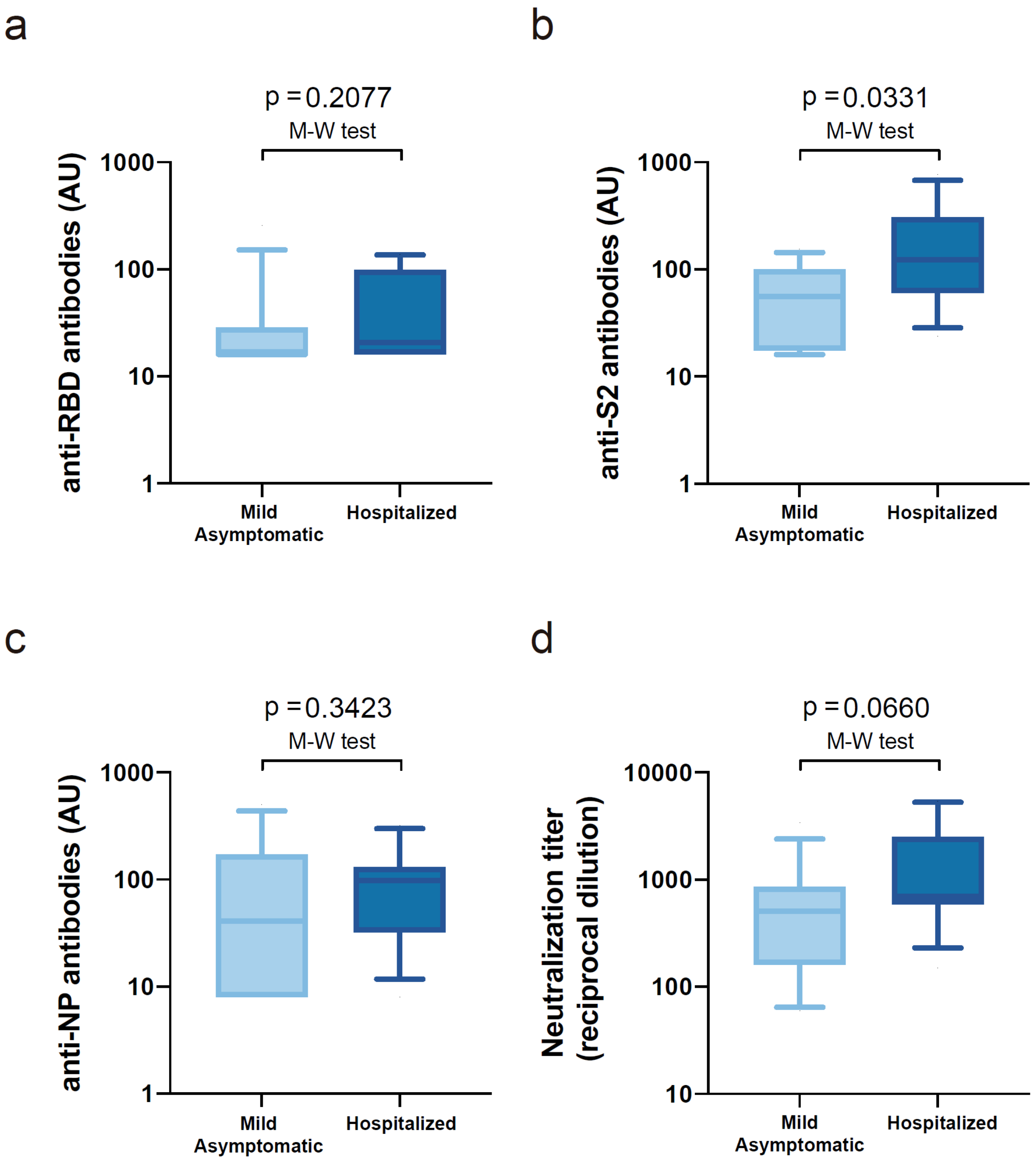


**Figure S2. Antibody titers at the end of the follow-up period.** Antibody titers of the last measure for IgG against the receptor binding domain (RBD) (**a**), S2 (**b**), and nucleoprotein (NP) (**c**) on a subset of individuals with largest follow-up (n=14 for mild/asymptomatic and n=14 for hospitalized). Panel **d** shows the neutralizing activity of the same subset of individuals at the end of the follow-up period. Boxes show the median and the interquartile range, and bars the $10^{th}$ and $90^{th}$ percentiles. Severity groups (i.e., mild/asymptomatic and hospitalized) were compared using the Mann-Whitney test.
